# Pregnancy lab test dynamics resemble rejuvenation of some organs and aging of others

**DOI:** 10.1101/2025.02.24.639848

**Authors:** Ron Moran, Glen Pridham, Yoel Toledano, Uri Alon

## Abstract

Aging and pregnancy both involve changes in many physiological systems. Some of these changes are similar, leading to suggestions that pregnancy may be a model for aging. Recent studies using DNA methylation clocks showed apparent aging during gestation which resolves postpartum. Since aging and pregnancy are complex, it is important to compare them in terms of many physiological parameters and at many time points. Here, we analyzed cross-sectional data on 62 lab tests at weekly resolution in 300,000 pregnancies and 1.4 million nonpregnant females aged 20-80. We trained a regression model to predict age from lab tests. Apparent age dropped by 8 years in early pregnancy, rose by 30 years towards delivery, and recovered postpartum. Certain systems exhibited rejuvenation, with opposite trends in pregnancy and aging, including renal, iron, and most liver tests. Others, such as coagulation, thyroid, muscle, and metabolic systems, showed apparent aging. Some systems displayed mixed trends. Notably, in the systems that showed apparent aging, the physiological mechanisms for the changes differed between pregnancy and aging. Pregnancy complications led to an additional apparent aging of 4-8 years. We conclude that pregnancy has features of rejuvenation for some systems, but its aging-like features involve different mechanisms than true aging. Gestational rejuvenation-like mechanisms may offer clues for slowing aspects of biological aging.

## Introduction

Both pregnancy ^1,2^ and aging ^3,4^ are associated with stress on the body and involve changes in many physiological systems. Some of the changes in pregnancy resemble changes in aging, such as a shift in immunity towards innate immunity ^5,6^ and reduced red blood cell (RBC) counts ^1,7^. This has led to the suggestion that pregnancy may be a model for aging ^8,9^. For example, pregnancy and its complications involve inflammation, oxidative stress and telomere changes ^10–14^, and similar changes also occur in aging.

The relation of pregnancy to aging has recently gained attention due to aging clocks. Aging clocks are statistical models designed to predict biological age from molecular cues, where biological age is defined as the age at which the mean population has an equivalent state of health to the individual ^4^. A major advance was the advent of clocks based on DNA methylation ^15,16^ in 2013 - several hundred methylation sites out of millions could accurately predict chronological age. Biological age acceleration was based on the residuals between true and predicted chronological age.

Next-generation clocks were trained directly on risk of morbidity and mortality, improving their specificity to biological age. These clocks were recently extended to the level of individual organs ^17–20^. Many clocks use additional molecular cues such as proteomics data ^21^ and blood tests ^22^. Using first- and second-generation clocks, a higher biological age was registered during pregnancy ^23,24^, which recovered after delivery.

If pregnancy and aging are related, knowledge may be transferred to treat aging related diseases; similarly, if pregnancy shows rejuvenation, it may inspire approaches for slowing aging. However, pregnancy and aging are complex and involve many systems. It is therefore worthwhile to make a detailed system-by-system comparison. Furthermore, the clock studies on pregnancy involve only one or two timepoints during gestation and one point postpartum. Higher temporal resolution can help resolve the dynamics of each trimester and the different stages of postpartum adaptation ^1^.

To compare aging and pregnancy, we study 62 laboratory (lab) blood tests comprising all major physiological systems from a nationwide dataset from the Clalit healthcare services ^25,26^. We use cross-sectional data from 313,000 pregnancies in weekly bins ^1^ and compare this to cross-sectional data from 1.4 million nonpregnant females at ages 20-89 ^25^. We develop a blood test aging clock and compare effective clock age per test and per system, each week from 60 weeks before delivery to 80 weeks after delivery, spanning preconception, gestation and postpartum periods.

During pregnancy, apparent lab-test age dropped by 8 years in the first trimester and then rose linearly, peaking at 30 apparent years older than baseline. After delivery, apparent age recovered gradually over a year postpartum. Several organ systems showed apparent rejuvenation - opposite changes in aging and gestation - including renal, iron and most liver tests. Other systems showed apparent aging - changes in the same direction in aging and gestation - such as coagulation, thyroid, muscle and metabolic tests. A few systems showed mixed directions compared to aging. Complications of pregnancy - preeclampsia and gestational diabetes - showed aging-like changes of 4-8 years compared to healthy pregnancies. The physiological mechanism underlying the test changes differed between pregnancy and aging.

## Results

### Lab-test ‘age’ drops during the first trimester, rises in the second and third, and resets over months postpartum

To evaluate the effective lab-test age during each week of pregnancy and postpartum, we developed a lab-test aging clock (Methods), based on linear regression between binned lab test mean values and chronological age. Reference values for healthy females are from LabNorm ^25^. We call this regression model LabAge.

LabAge predicts chronological age with r=0.87, p-value <0.001, similar to other aging clocks that use blood tests ^22,27^. The goal here is to provide a measure in years for the apparent aging and rejuvenation of tests during pregnancy and postpartum. The lab tests that contributed the most to LabAge are urea, systolic blood pressure, the glycemic test HbA1c, LDL cholesterol, creatinine, red blood cell distribution width (RDW-standard deviation) and Albumin; all rise with age except albumin which decreases. The weights for each test are shown in Fig 1.

**Fig. 1.**
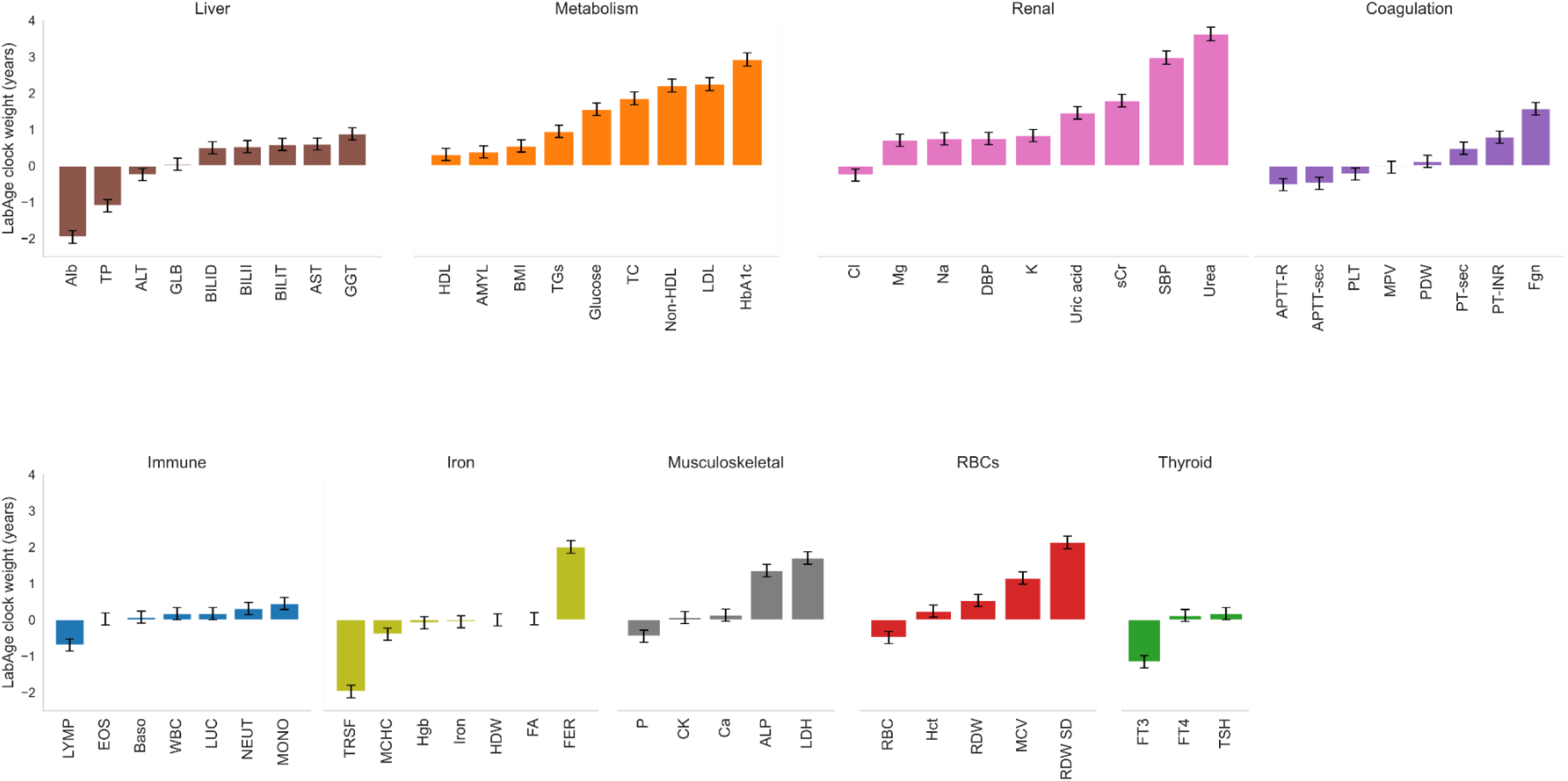
Weights for the lab test regression clock (LabAge). Tests are grouped according to system. Folic acid (FA) and hemoglobin related RBC tests are grouped with iron tests. sCr is serum creatinine, TC is total cholesterol. Full test names are in table S1 (supplementary information). Weights are for LabAge, a regression model trained on a z-scored dataset for comparability.

We applied LabAge to pregnancy cross-sectional data binned by weeks. For each week we used the mean test value of all participants tested in that week. The dataset ^1^ spans a period starting from 60 weeks before delivery to 80 weeks after delivery. Each lab test is evaluated in terms of quantiles of a reference population of healthy age-matched non-pregnant females ^1^ (see Methods).

The LabAge clock provides an age for each week of pregnancy according to the lab test data. Apparent lab-test age drops in the first trimester by about eight years, then rises linearly with gestation week peaking at delivery with an apparent aging of 30 years above preconception (Fig 2). LabAge then recovers halfway at ten weeks postpartum, and another halfway at 60 weeks postpartum. The postpartum period thus shows fast (10 weeks) and slow (year) recovery phases.

**Fig. 2.**
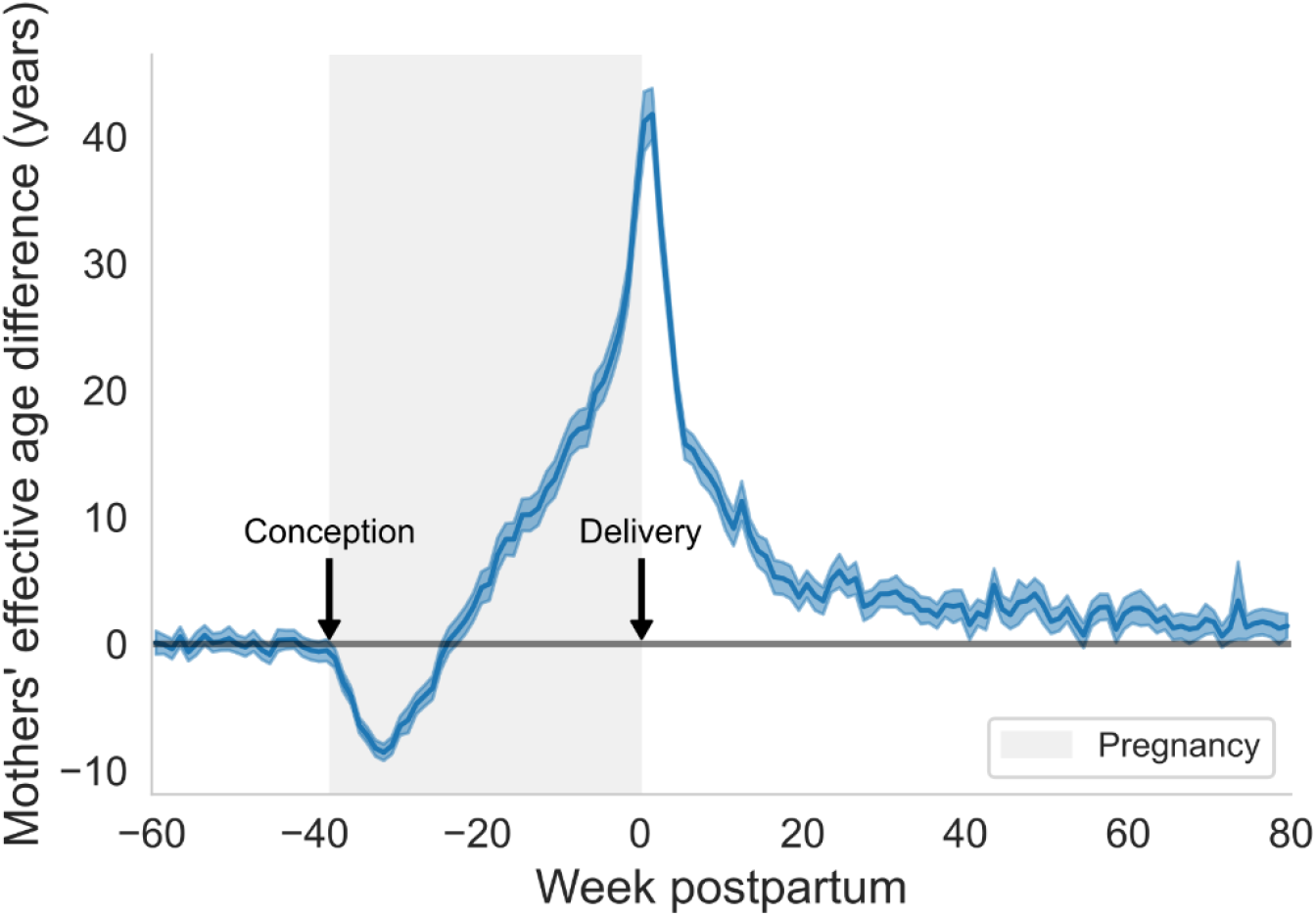
Mothers’ mean lab test age drops in trimester one, rises towards delivery and recovers postpartum. LabAge at each week of gestation and postpartum where age 0 is baseline clock age at 60 weeks before delivery. Gestation is in gray background; delivery is at t=0. Errors (shaded) are 95% confidence interval (CI) from resampling (see Methods). Outliers were corrected using the empirical distribution per weekly bin (see Methods).

### Some systems show apparent rejuvenation, while others show apparent aging

We calculated an effective age for each test using the regression model. In Fig 3, the test ages are grouped by physiological systems. We computed the effective age change for each system using these grouped tests (black curves in Fig 3). Most systems have one or two tests that contribute most strongly to its effective age.

**Fig. 3.**
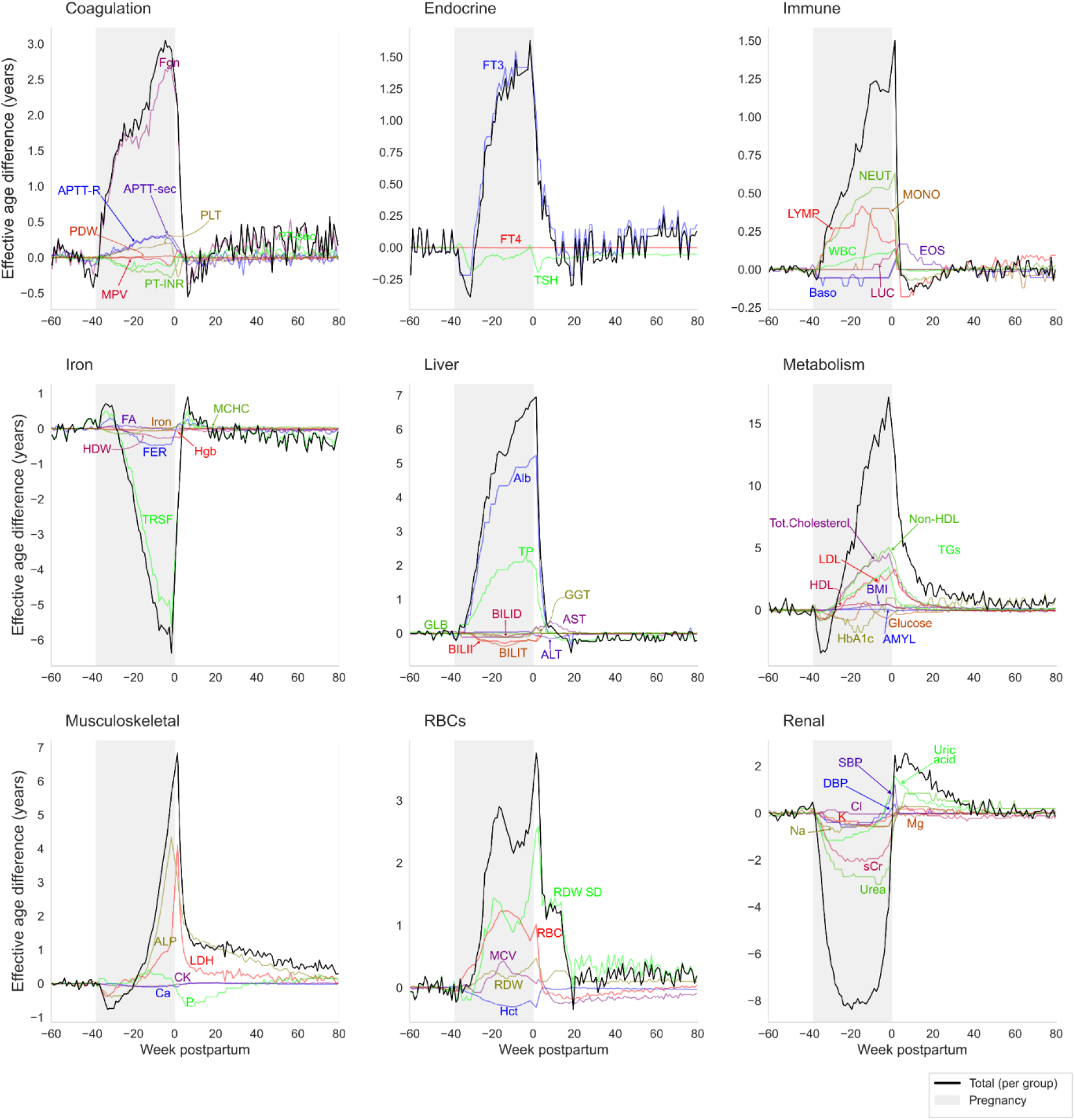
Apparent age per week of pregnancy and postpartum by test and system. Thick black lines are the total change per system, and individual test age traces are labeled and in colors. Gestation is in gray background, delivery is at t=0. Total changes sum up to the effective age difference in Fig. 2.

Based on the lab tests, the renal and iron (including hemoglobin) tests show the greatest apparent rejuvenation. Conversely, other systems show apparent aging. The largest aging effects are in the metabolic, hepatic and musculoskeletal systems, followed by immune, RBC, coagulation and thyroid tests. In the first trimester, some changes are opposite to the second and third in several systems like coagulation, iron, muscle and metabolism.

After delivery many systems recover within weeks. In the postpartum period. Other systems, like the immune system, show an undershoot, whereas the renal tests display a sizable overshoot. The slowest postpartum recovery is in the musculoskeletal and metabolic tests. We provide more details on each system next.

### Renal and iron lab-test changes in pregnancy are opposite to those in aging - apparent rejuvenation

To further compare the effects of aging and pregnancy, we compare in Fig 4 changes in aging to changes in pregnancy test by test. The y axis is the maximal quantile change during gestation, and the x axis is the quantile change between age 20 and 80. Apparent aging occurs along the first and third quadrants, and apparent rejuvenation along second and fourth quadrants.

**Fig. 4.**
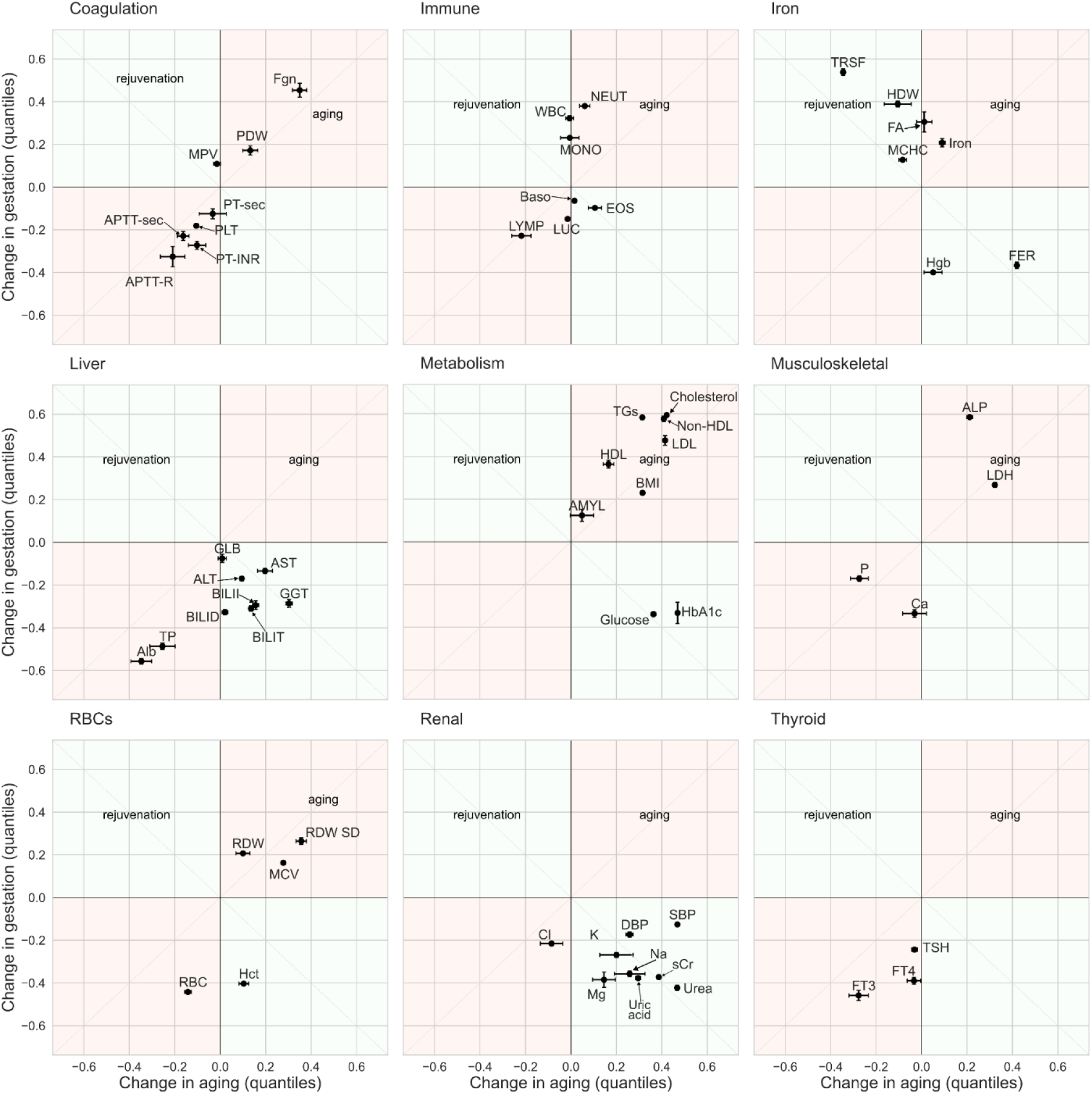
Changes in aging versus pregnancy show apparent rejuvenation or aging in different tests and systems. Maximal quantile change (max-min) in gestation versus change between age 20 and 80 (see Methods), in quantiles of the reference population (healthy nonpregnant females). Quadrants 1 and 3 (pink) denote changes in the same direction and are labeled ‘aging’, and quadrants 2 and 4 (green) denote changes in opposite directions and are denoted ‘rejuvenation’. Error bars are 2 standard deviations (see Methods).

Renal tests including electrolytes, urea and creatinine showed sizable changes in both aging and gestation, but these changes had opposite directions as shown in Fig 4. This apparent rejuvenation during gestation reflects the increase in glomerular filtration rate (GFR) during pregnancy of 30% ^28^, as opposed to the 40% decrease in GFR over aging ^29^.

Tests related to iron and hemoglobin also showed opposite directionality in pregnancy and aging. Hemoglobin and ferritin dropped in pregnancy and rose in aging, and transferrin rose in pregnancy and dropped in aging. This reflects aspects of mild anemia found in pregnancy ^1,30,31^ and excess iron found in aging ^32^. Folic acid tests are affected by pregnancy supplements and rise in pregnancy ^33^ but are unaffected in aging.

### Liver test changes in pregnancy are mostly opposite to those in aging - indicating apparent rejuvenation - except for albumin

We find that liver tests that represent hepatocellular dysfunction, e.g. ALT and AST, dropped in pregnancy and rose in aging (Fig 4).

Two other liver tests showed changes in the same direction as aging, dropping with age and gestation - albumin and total protein (the two tests are strongly correlated in general). However, the origin for this decline is different in aging and pregnancy. During pregnancy, the drop reflects increased hemodilution due to the 50% rise in blood volume ^34^, whereas in aging the drop is due to decline in liver albumin production at old age ^35,36^.

### Coagulation, thyroid, muscle and metabolic tests show apparent aging during gestation

Coagulation tests indicate a hypercoagulative state in both aging and gestation (Fig 4). Hypercoagulation in gestation is thought to be protective ^37^ since delivery is associated with blood loss. Aging is also associated with hypercoagulation. However, hypercoagulation at old age is detrimental, and is linked to endothelial dysfunction^38^ and to the inflammatory state in old age known as inflammaging ^39^.

Many metabolic tests show increased LabAge. lipid tests and BMI increase both in gestation and aging, including serum LDL, triglycerides and total cholesterol (Fig 4). Pregnancy is associated with weight gain and increased BMI, due mainly to elevated blood and amniotic fluid volume and the growing fetus ^40^. In contrast, in aging the rise in BMI is driven mostly by elevated fat mass^41–43^

We find that glycemic tests show complex dynamics during gestation. This is due in part to insulin sensitivity in the first trimester and insulin resistance in the second and third. The maximal trend is negative. Hence, it is opposite to the rise of glycemic tests in aging.

The apparent rejuvenation of glycemic tests is driven ^44^ by placental hormones and by the demands of the embryo. In aging, elevated glucose is driven in part by insulin resistance due to inflammaging, low activity and increased visceral fat.

The thyroid tests in pregnancy include a suppression of serum thyroid stimulating hormone (TSH) (Fig 4), partly due to the TSH-like effects of the placental hormone human chorionic gonadotropin (hCG) in the first trimester ^45^.

A reduction of free Thyroxine ( FT4) is also noticeable. Both pregnancy and aging show a drop in free triiodothyronine (FT3). During gestation, this is due to increased renal iodine clearance and higher expression of thyroid binding globulin (TBG) ^46^. On the other hand, aging is associated with a reduced metabolic demand which affects the deiodination of T4 to T3. Moreover, the euthyroid sick syndrome related to inflammation and chronic illness is more prevalent in old age ^47–49^.

The musculoskeletal tests generally show changes aligned with aging. One of the lab tests that changes most in pregnancy, serum alkaline phosphatase (ALP), rises also in aging, in part due to the expression of a placental variant of the ALP enzyme ^50^.

### Red blood cell dynamics and immune system tests present mixed similarity with aging

Immune and red blood cell tests display a more complex picture (Fig 4). More immune-related blood tests vary in pregnancy than in aging. The test that varies most in aging is a reduction in lymphocytes. It shifts in a similar direction in pregnancy ^51^. The mechanisms in both states are distinct. In pregnancy, the adaptive immune system is hormonally redirected to avoid attacking the embryonic tissues ^6,52^. In aging, thymic involution and other alterations drive a reduction in lymphocyte counts ^53,54^.

Generally, red blood cell tests shift during gestation and aging in the same direction. Specifically, RBC count drops while mean corpuscular volume (MCV) and red blood cells distribution width (coefficient of variation, RDW) both rise ^7,51,55^. However, the hematocrit shows opposite effects, rising in aging and dropping in pregnancy. The latter effect is probably due to hemodilution ^2,56^.

### Pregnancy complications display lab test aging relative to a healthy pregnancy

We compared the LabAge of healthy pregnancies to three of the most common severe pregnancy complications: preeclampsia, gestational diabetes and postpartum hemorrhage (PPH) (Fig 5). Preeclampsia involves high blood pressure, organ damage and risk of seizures, gestational diabetes is hyperglycemia without history of a previous diabetes, and PPH involves severe bleeding which can be life threatening. Data is from Bar et al^1^.

**Fig. 5.**
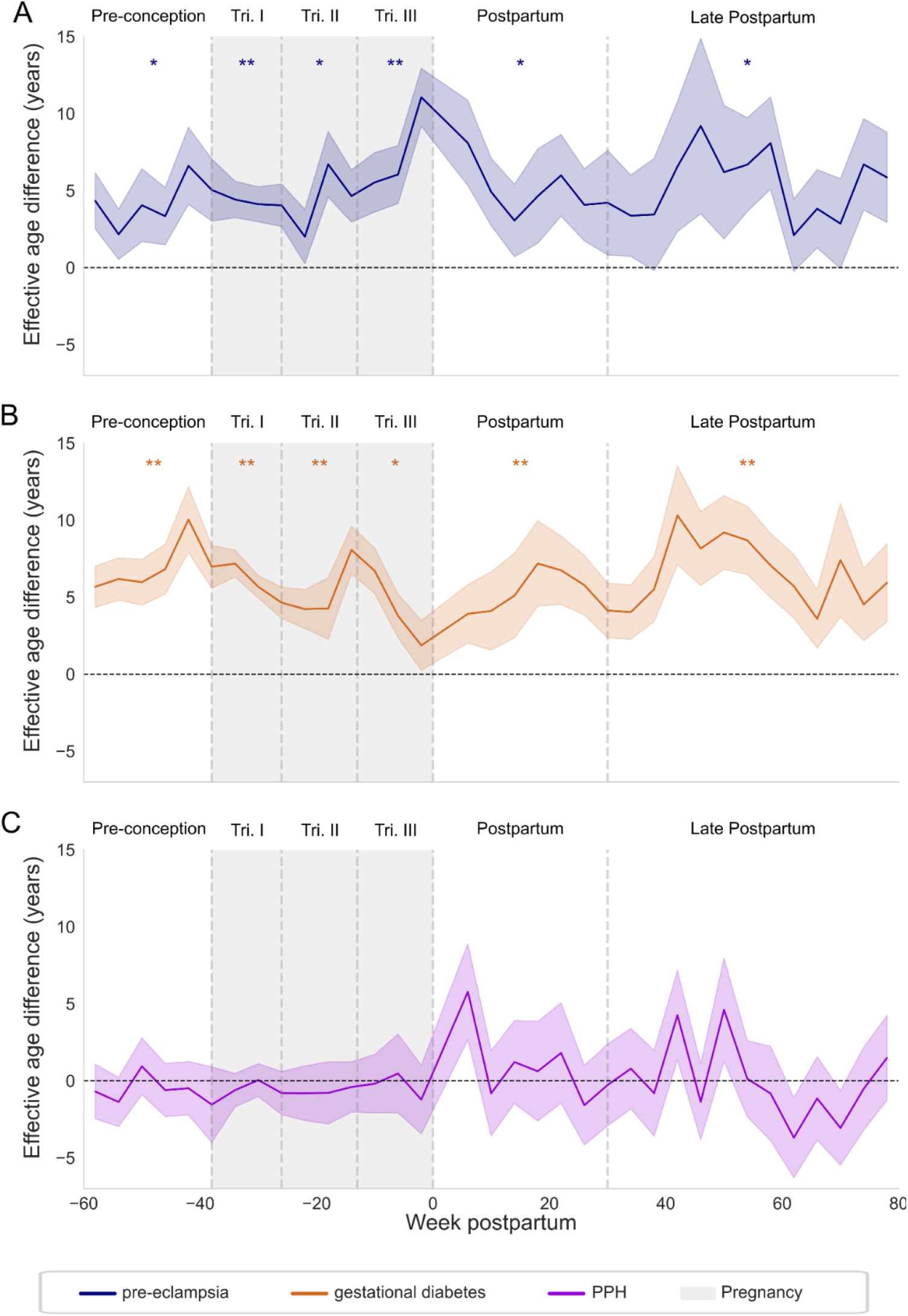
Pregnancies complicated by preeclampsia and gestational diabetes shows higher apparent age than healthy pregnancies. A. preeclampsia n=5643 pregnancies b. gestational diabetes, n=7244 c. PPH n=4574. Data is grouped by 4 week intervals. Statistical significance (one way z test) for positive age was computed for preconception, for each trimester and for early and late postpartum with dashed lines demarcating these periods. Significance is denoted by *p<0.05, ** p<0.01 with multiple hypothesis testing correction (Benjamini Hochberg). Errors (shaded) are 95% CI from resampling (see Methods).

Preeclampsia and gestational diabetes cohorts showed LabAge that is about 5 years older than healthy pregnancy. Preeclampsia LabAge seems to rise further in the third trimester.

The increased lab test age appears also before gestation, because the population of women at risk differs from the general population. Women with both complications exhibit metabolic changes and inflammation markers which also characterize aging ^8^. For example, some alterations include elevated glucose, serum LDL cholesterol, blood pressure and BMI.

PPH did not show a significant aging effect except right after delivery. Since this complication occurs at delivery it may not involve physiological changes as pronounced as the other two complications, which are diagnosed during gestation ^57–59^.

When comparing the changes test by test, pregnancies complicated by preeclampsia and gestational diabetes differs from healthy pregnancies in a direction that aligns with aging (Fig S1, SI).

## Discussion

We compared the weekly changes in 62 lab tests during pregnancy to their changes during aging, using cross-sectional data from 313,000 pregnancies and 1.4 million nonpregnant females at ages 20-89. The results highlight that while pregnancy and aging share some physiological changes, they are nevertheless unique processes involving different biological mechanisms. Apparent lab-test age during pregnancy dropped by 8 years in the first trimester and rose by 30 years towards delivery, recovering gradually over postpartum on a timescale of weeks with a slow recovery tail over a year. Several systems showed apparent rejuvenation including renal, iron and most liver tests. Other systems, such as coagulation, thyroid, muscle and metabolic tests, showed apparent aging. A few systems showed mixed directions in pregnancy vs. aging. Complications of pregnancy showed aging-like shifts of about 5 years compared to healthy pregnancies.

The rejuvenation of renal tests stems primarily from the rise in glomerular filtration rate (GFR) during gestation, by a factor of about 30%, starting in the first trimester. This rise is driven by systemic vasodilation, plasma volume expansion, caused by placental hormonal influences including relaxin, progesterone, and estrogen. Collectively, these hormones reduce vascular resistance and enhance renal perfusion ^28,60^. In aging, GFR gradually declines due to reduced cardiac output and accumulating kidney damage causing lab tests to shift in the opposite direction.

The apparent rejuvenation of iron related tests stems from increased iron storage during aging ^32^, in contrast to an increased demand for iron during gestation ^31^.

Thus, pregnancy shows aspects of physiological rejuvenation for some organ systems which may be considered as reverse aging-such as increased kidney filtration rate and cardiac output throughout gestation and increased insulin sensitivity in the first trimester. Many of these effects occur already in the first weeks of the first trimester, and are caused primarily by placental hormone production including hPL, estrogen, progesterone and relaxin. Other changes such as increased blood volume and adiposity occur later in gestation. Peak rejuvenation is seen around the end of the first trimester and the beginning of the second trimester.

These effects might inspire approaches to mimic rejuvenation in aged individuals. Such approaches would not be as straightforward as administering the placental hormones at pregnancy doses since they would cause severe side effects.

In contrast to rejuvenation, other organ systems show lab test changes that resemble aging. However, the mechanistic reasons for the aging-like changes in pregnancy are usually different from true aging. Many of the changes in pregnancy are due to uncompensated hemodilution and the metabolic demands of the fetus in trimesters 2 and 3. For example, blood volume grows by about 50% over gestation causing hemodilution^34^, and thus tests for serum proteins like albumin drop during pregnancy. Albumin also drops during aging, but for different reasons, primarily a decline in its liver production rate due in part to downregulation due to chronic inflammation as a negative acute phase response^53,61,62^. Another major physiological difference is the involvement of placental hormones which are absent in aging. These hormones In trimesters 2 and 3 cause extensive changes in lipids, glucose, coagulation, musculoskeletal, immune tests and others.

A transient rise in methylation clock age is seen in pregnancy, and a similar rise is also seen in surgery, COVID-19 and other conditions ^23^. Existing methylation clocks may thus measure not biological age per se, but rather physiological stress. One potential advance would be to develop accurate clocks for biological age based on molecular mechanisms specific to aging rather than general lab tests or methylation. Thus, it may be possible to avoid being confounded by extrinsic, physiological and stochastic ^63,64^ responses that are not related to aging.

We find that pregnancy complications such as preeclampsia and gestational diabetes differ from healthy pregnancy in a way that aligns with changes in aging. They add an apparent 5 years to the LabAge lab test clock. This apparent aging exists before pregnancy and does not change significantly with gestation and postpartum, suggesting that it is a characteristic of the population susceptible to these pregnancy complications. In contrast, PPH, which manifests after delivery did not significantly alter lab test age compared to healthy pregnancy except for the weeks after delivery. High LabAge may thus be an indicator of risk for certain complications; it would be of interest to construct classifiers to predict pregnancy complications, but the cross-sectional nature of the data does not allow us to do so in the present study.

We conclude that cross-sectional data with high temporal resolution helps to resolve the complexity of physiological change in pregnancy, with some systems showing rejuvenation and others apparent aging. Opposite effects were documented in the first trimester compared with the second and third trimesters. Postpartum recovery occurred rapidly within weeks and then more slowly over about a year. The rejuvenating features of pregnancy may offer clues for slowing aspects of biological aging.

## Methods

### Study population (pregnancy dataset)

The study population consists of Clalit healthcare members who were pregnant in the years 2003-2020 aged [20,35]. Only healthy pregnancies were considered except as noted. Due to privacy considerations, the analysis was on summary statistics from participants who took the lab tests in each weekly bin. Individualized data are not available. For more information see Bar et al ^1^.

### Reference Population

Lab test distributions of healthy, non-pregnant female populations parameterized by age are from LabNorm ^25^.

### Linear regression

The LabAge clock model was based on ordinary least squares regression using the python module statsmodels.api.OLS. CI on the parameters was computed using statsmodels.regression.linear_model.RegressionResults.conf_int with 95%. We trained the regression model on reference data from LabNorm. Each test has reference values for ages 20 to 89 for females. Pearson r was reported between predicted and sampled age. All tests used are shown in table S1 (supplementary information).

### Maximum quantile difference in pregnancy

To estimate max difference and its error bars, we constructed a distribution parameterized by the mean quantile score and standard deviation for each gestational week (measured by 0 as delivery) indexed by and lab test indexed by *t*:

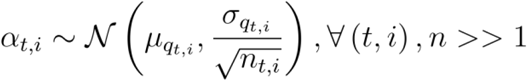

We sampled over the gestation weeks *α*_*t*,*i*_,*∀*_*i*_ ∈[−38,0]. We calculated the maximal difference for each such sampling indexed by *j*:

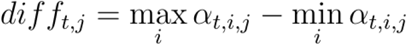

To resample, we repeated for 1000 *j*’s and reported the mean and CI of 2 standard deviations ∼97%.

### Quantile difference in aging

To compute the quantile difference between age 80 and 20, we estimated the age 80 value by the median LabNorm values at the age range ∈[75,84] for healthy females. If there was a significant change within this age range, as defined by regression p value <0.05, we used the residuals of the regression at age 80. We then transform these values into quantiles relative to the reference population at age 20 and subtract 0.5 which is the mean quantile score by definition. CI were 2 standard deviations ∼97%.

### CI for LabAge

Mean values per gestational week and test were resampled 1000 times as above. Parameters from the regression were sampled assuming normality 1000 times for each weekly bin *i*. We report the resampling mean and the CI of 1.96 standard deviations ∼ 95%. In Fig 5, The weekly bin of delivery was excluded due to high variance across all lab tests.

### Outlier correction

Some weekly bins in some tests have strong outliers shifting the mean and standard deviation. For test *t* and weekly bin *i*, we defined outliers by standard deviation that exceeds 10 times the difference between the ∼ 95^th^ and the 5^th^ percentiles:

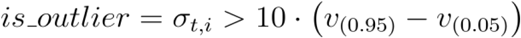

We corrected such outliers using linear interpolation of the distribution of the adjacent weekly bins. Denoting the PDF as 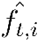:

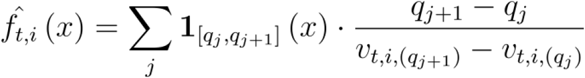

Extrapolating left to *q* _0_= 0 and right to 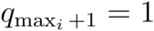 and vanishing outside. We calculated the mean and standard deviation by definition for continuous CDF 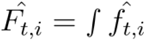. The uncorrected prediction in Fig 2 is given in Fig S2 (supplementary information).

### p-values in the pregnancy complication prediction (one-way z-test)

LabAge of the reference was subtracted from LabAge of each complication. For each gestational period (preconception, trimesters, early and late postpartum) we pooled all

LabAges and calculated: 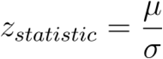. We reported the p-value as:1−Φ(*z*_*statistic*_). Controlling for false discovery rate (FDR) used Benjamini Hochberg for the 6 p-values.

## Acknowledgments

We thank all members of our lab and Ido Solt, Amos Tanay and Neta Mendelsohn for discussions. We thank Gabi Barabash and Ran Balicer for the Clalit−Weizmann collaboration. Data acquisition was approved by the Clalit Helsinki Committee RMC-1059-20.

## Author contributions

Conceptualization: UA, RM, GP

Methodology: RM, GP, UA

Formal Analysis: RM, GP

Funding acquisition: UA

Visualization: RM, UA

Data Curation: YT

Supervision: UA

Software: RM

Writing – Original Draft: UA, RM

## Data and materials availability

All data needed to evaluate the conclusions in the paper are present in the paper and/or the Supplementary Materials. The source code and data used to perform the analysis is available at the GitHub repository as of the date of publication. The repository is open for public use:

**https://github.com/AlonLabWIS/PregAging**

