## Supplementary Information for "Pregnancy lab test dynamics resemble rejuvenation of some organs and aging of others"

This supplementary includes:
Figures (fig S1, fig S2)
Tables (table S1)

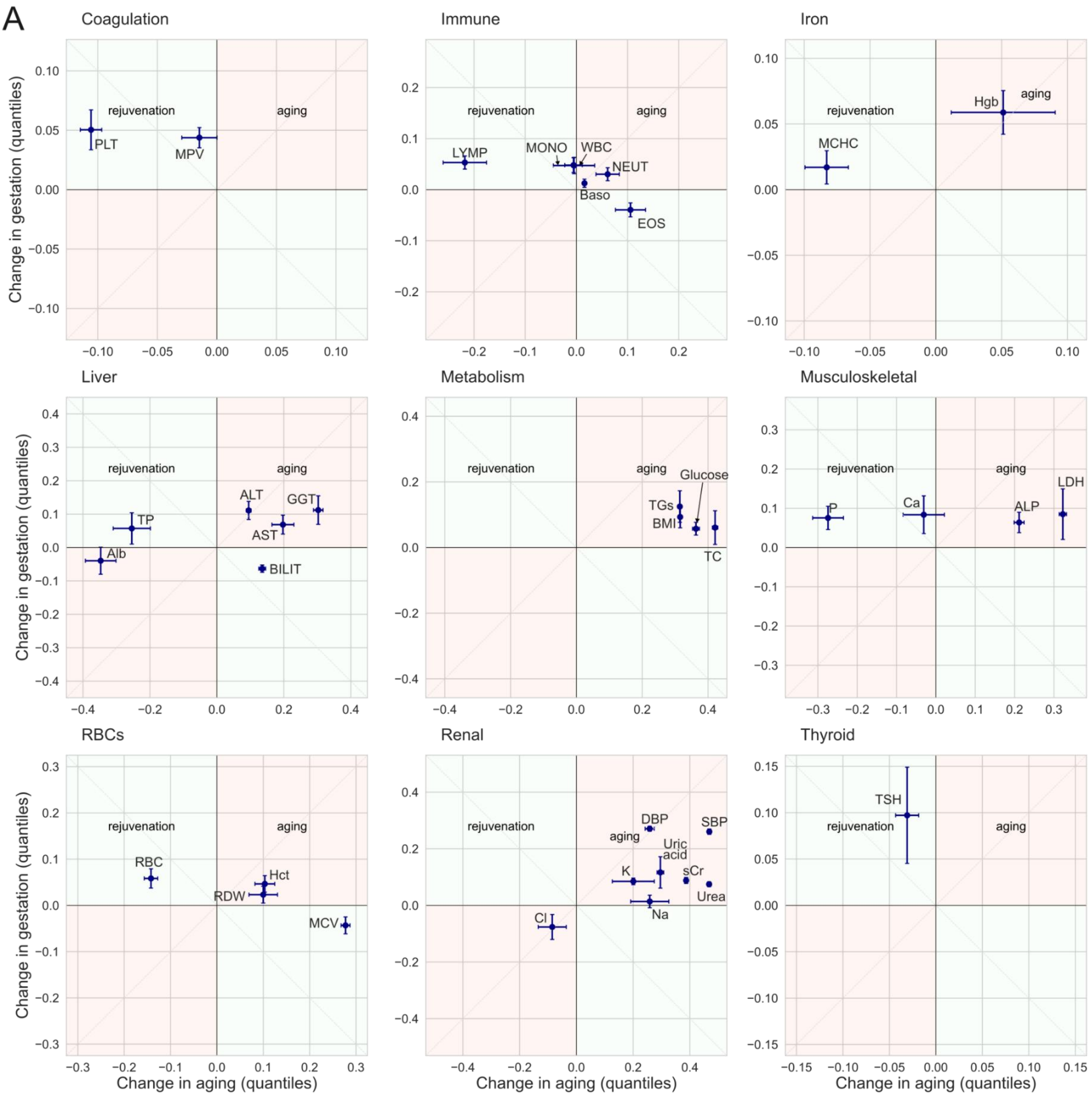
A
Coagulation
Immune
Iron
Liver
Metabolism
Musculoskeletal
RBCs
Renal
Thyroid
rejuvenation
aging
PLT
MPV
LYMP
MONO
WBC
NEUT
Baso
EOS
Hgb
MCHC
ALT
GGT
TP
AST
Alb
BILIT
Glucose
TGs
BMI
TC
LDH
P
Ca
ALP
RBC
Hct
RDW
MCV
DBP
SBP
Uric acid
K
sCr
Urea
Na
Cl
TSH
Change in gestation (quantiles)
Change in aging (quantiles)

B

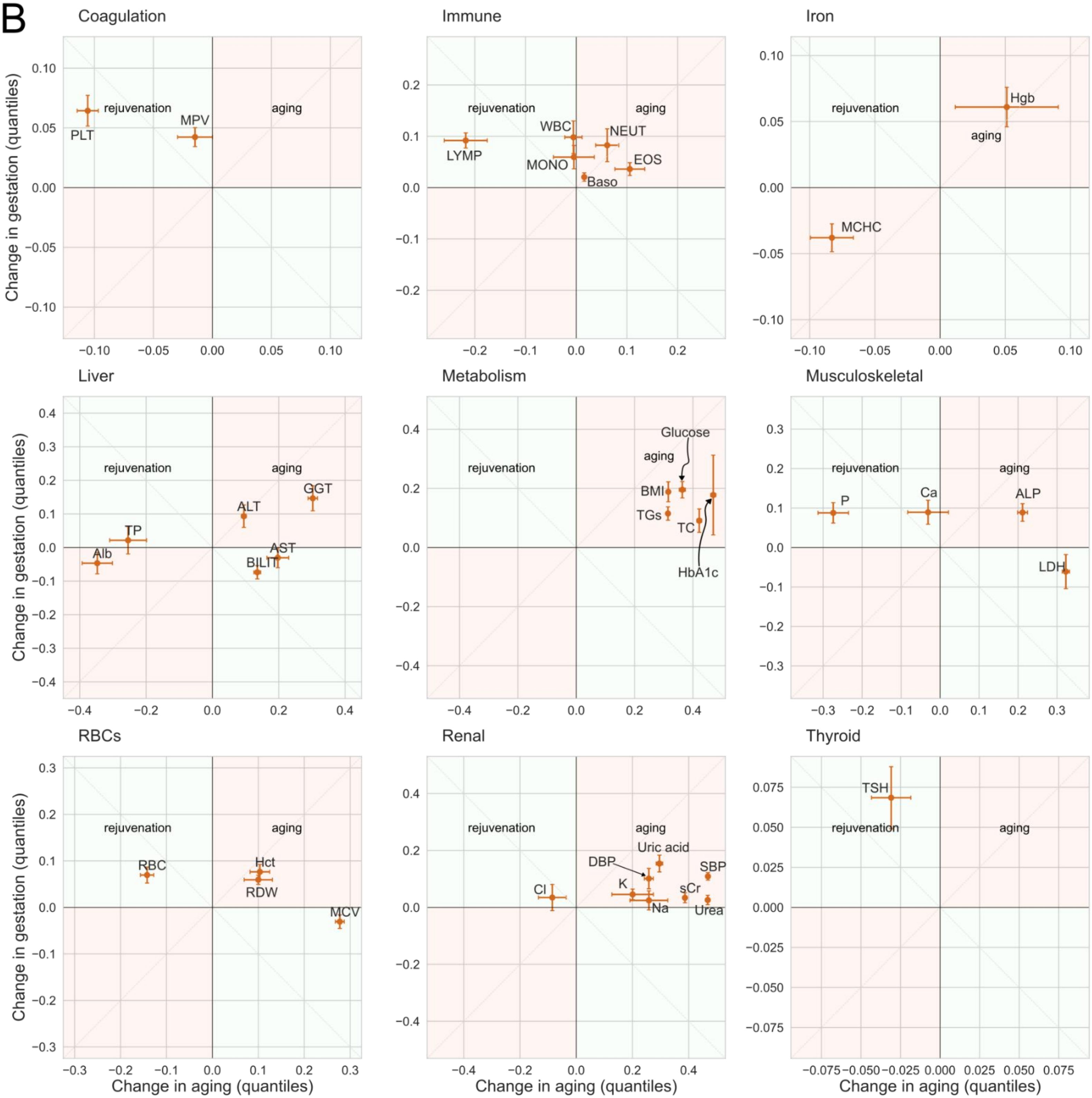
Coagulation
Immune
Iron
Liver
Metabolism
Musculoskeletal
RBCs
Renal
Thyroid
rejuvenation
aging
PLT
MPV
LYMP
WBC
MONO
NEUT
EOS
Baso
Hgb
MCHC
GGT
ALT
TP
Alb
AST
BILIT
Glucose
BMI
TGs
TC
HbA1c
P
Ca
ALP
LDH
RBC
Hct
RDW
MCV
Uric acid
DBP
SBP
K
Cl
sCr
Na
Urea
TSH
Change in gestation (quantiles)
Change in aging (quantiles)

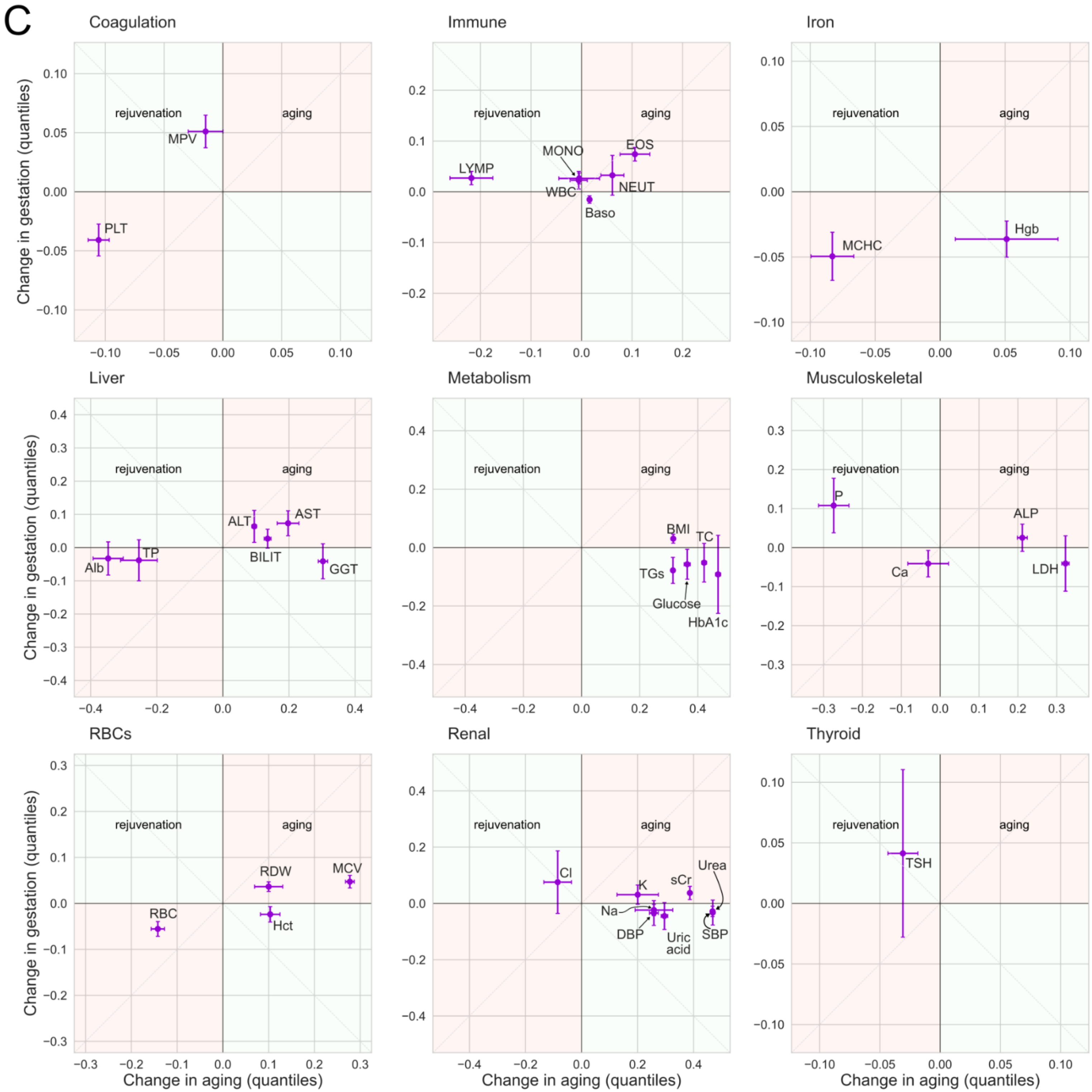


**Fig S1 - Comparison of lab test quantile changes in pregnancy complications and aging.** Shown are maximum quantile differences between healthy pregnancies and (a) preeclampsia, (b) generational diabetes and, (c) PPH versus changes in aging.

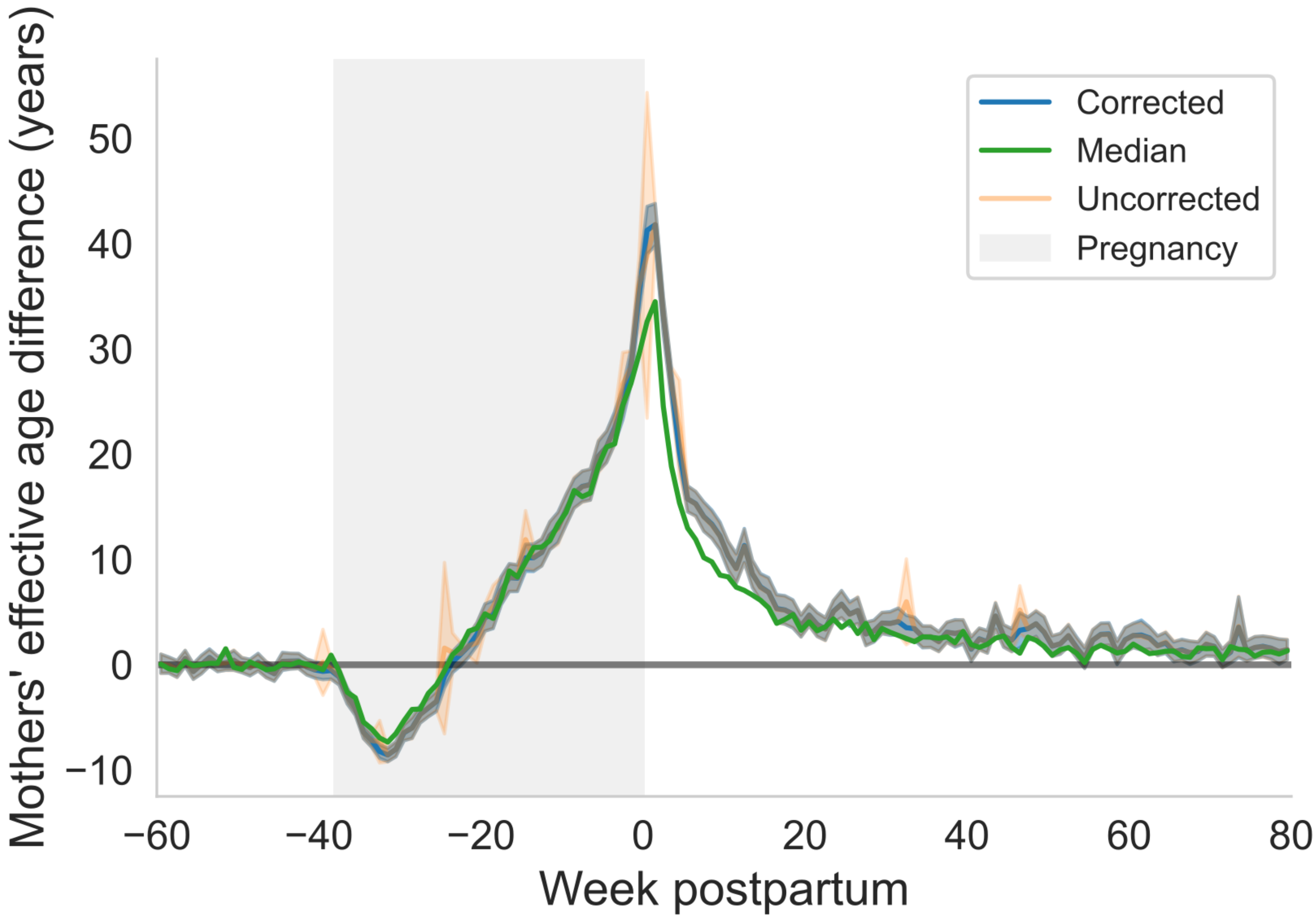


**Fig S2. Outlier-corrected and uncorrected prediction by LabAge.** Uncorrected values have larger errors 25 weeks before delivery and at delivery. As a reference, we generated a LabAge prediction based on median values for each test and weekly bin instead of mean values used in the corrected and uncorrected prediction.

| Name | Full name | Abbreviation | Group | Units |
|---|---|---|---|---|
| Albumin | ALBUMIN | Alb | Liver | g/dL |
| ALT | ALT_Alanine_aminotransferase_GPT | ALT | Liver | units/L |
| Amylase (*) | AMYLASE_BLOOD | AMYL | Metabolism | units/L |
| APTT-R (*) | APTT_R | APTT-R | Coagulation | |
| APTT-sec (*) | APTT_sec | APTT-sec | Coagulation | seconds |
| AST | AST_Aspartate_aminotransferase_GOT | AST | Liver | units/L |
| Basophiles | BASOPHILES_abs | Baso | Immune | K-cells/μL |
| Bilirubin(dir) (*) | BILIRUBIN_DIRECT | BILID | Liver | mg/dL |
| Bilirubin(indir) (*) | BILIRUBIN_INDIRECT | BILII | Liver | mg/dL |
| Bilirubin(tot) | BILIRUBIN_TOTAL | BILIT | Liver | mg/dL |
| BMI | BMI | BMI | Metabolism | $kg/m^2$ |
| BP (Diastolic) | BPd | DBP | Renal | mmHg |
| BP (Systolic) | BPs | SBP | Renal | mmHg |
| Calcium | CALCIUM_BLOOD | Ca | Musculoskeletal | mg/dL |
| Cholesterol | CHOLESTEROL | TC | Metabolism | mg/dL |
| HDL (*) | CHOLESTEROL_HDL | HDL | Metabolism | mg/dL |
| LDL (*) | CHOLESTEROL_LDL | LDL | Metabolism | mg/dL |
| Creatine kinase | CK_CREAT | CK | Musculoskeletal | IU/L |
| Chloride | Cl | Cl | Renal | |
| Creatinine (blood) | CREATININE_BLOOD | SCr | Renal | mg/dL |
| Eosinophils | EOS | EOS | Immune | K-cells/μL |
| Ferritin (*) | FERRITIN | FER | RBCs | ng/mL |
| Fibrinogen (*) | FIBRINOGEN | Fgn | Coagulation | mg/dL |
| Folic acid (*) | FOLIC_ACID | FA | RBCs | nmol/L<ff> |
| GGT | GAMMA_GLUTAMYL_TRANSPEPTIDASE | GGT | Liver | IU/L |
| Globulin (*) | GLOBULIN | GLB | Liver | g/dL |
| Glucose† | GLUCOSE_BLOOD | Glucose | Metabolism | mg/dL |
| Hematocrit | HCT | Hct | RBCs | percent |
| HDW (*) | HDW | HDW | RBCs | |
| HbA1c | HEMOGLOBIN_A1C_CALCULATED | HbA1c | Metabolism | percent |
| Hemoglobin | HGB | Hgb | RBCs | g/dL |
| Iron (*) | IRON | Iron | RBCs | microg/dL |
| Potassium | K | K | Renal | mEq/L |
| LDH | LACTIC_DEHYDROGENASE_LDH__BLOOD | LDH | Musculoskeletal | units/L |
| Large unstained cells (*) | LUC | LUC | Immune | |
| Lymphocytes | LYMP | LYMP | Immune | K-cells/μL |

| | | | | |
|---|---|---|---|---|
| Magnesium (*) | MAGNESIUM_BLOOD | Mg | Renal | mEq/L |
| MCHC | MCHC | MCHC | RBCs | g/dL |
| MCV | MCV | MCV | RBCs | pg/cell |
| Monocytes | MONO | MONO | Immune | K-cells/μL |
| MPV | MPV | MPV | Coagulation | fL |
| Sodium | Na | Na | Renal | mEq/L |
| Neutrophils | NEUT | NEUT | Immune | 1000/microL |
| Non-HDL Cholesterol (*) | NON_HDL_CHOLESTEROL | Non-HDL | Metabolism | mg/dL |
| PDW (*) | PDW | PDW | Coagulation | |
| Alkaline Phosphatase | PHOSPHATASE_ALKALINE | ALP | Musculoskeletal | units/L |
| Phosphorus | PHOSPHORUS_BLOOD | P | Musculoskeletal | mg/dL |
| Platelets | PLT | PLT | Coagulation | K/microL |
| Total Protein | PROTEIN_TOTAL_BLOOD | TP | Liver | g/dL |
| PT INR (*) | PT_INR | PT-INR | Coagulation | g/dL |
| PT sec (*) | PT_SEC | PT-sec | Coagulation | seconds |
| RBC | RBC | RBC | RBCs | K-cells/μL |
| RDW | RDW | RDW | RBCs | |
| RDW SD (*) | RDW_SD | RDW SD | RBCs | |
| Free T3 (*) | T3_FREE | FT3 | Thyroid | pmol/L |
| Free T4 (*) | T4_FREE | FT4 | Thyroid | pmol/L |
| Transferrin (*) | TRANSFERRIN | TRF | RBCs | mg/dL |
| Triglycerides | TRIGLYCERIDES | TGs | Metabolism | mg/dL |
| TSH | TSH_THYROID_STIMULATING_HORMONE | TSH | Thyroid | |
| Urea | UREA_BLOOD | Urea | Renal | mg/dL |
| Uric acid | URIC_ACID_BLOOD | Uric acid | Renal | mg/dL |
| WBC | WBC | WBC | Immune | K-cells/μL |

† Glucose test – fasting glucose test. Instructions for the test include an 8 hour fast before drawing blood. No limitation on hydration.

**Table S1. Laboratory tests in the dataset**. Tests annotated with one asterisk (*) were excluded from regression analysis in the **“Complications of Pregnancy”** due to the low number of measurements in the cohort with complications.
